# On the quest for novelty in ecology

**DOI:** 10.1101/2023.02.27.530333

**Authors:** Gianluigi Ottaviani, Alejandro Martínez, Matteo Petit Bon, Stefano Mammola

## Abstract

The volume of scientific publications continues to grow, making it increasingly challenging for scholars to publish papers that capture readers’ attention. While making a truly significant discovery is one way to attract readership, another approach may involve tweaking the language to overemphasize the novelty of results. Using a dataset of 52,236 paper abstracts published between 1997 and 2017 in 17 ecological journals, we found that the relative frequency of novelty terms (e.g. *groundbreaking, innovative*) nearly doubled over time. All journals but one exhibited a positive trend in the use of novelty terms during the studied period. Conversely, we found no such trend for confirmatory terms (e.g. *confirm, replicated*). Importantly, only papers using novelty terms were associated with significantly higher citation counts and were more often published in journals with a higher impact factor. While increasing research opportunities are surely driving advances in ecology, the writing style of authors and the publishing habits of journals may better reflect the inherently confirmatory nature of ecological research. We call for an open discussion among researchers about the potential reasons and implications of this language-use and scientometrics issue.

## The recent rise in scientific production

“*Eureka!*”– yelled Archimedes when he solved a scientific problem that, among other things, would have cost him his life. This is only one of many tales of serendipitous discoveries that populate the history of science. A common thread in these narratives is the presence of a lonely genius who, perhaps in a stroke of luck or inspiration, succeeded in shedding light on the unknown (Conner, 2005). However, the reality behind these tales can be quite different (Foucault, 1969). Modern science is a systematized body of positive knowledge (Hoyningen-Huene, 2013), primarily built through a lengthy and steady accumulation of confirmatory work, only occasionally disrupted by game-changing discoveries that typically arise from anomalous results or observations (Darwin, 1859; Kuhn, 1962). Even after such discoveries, paradigms rarely shift abruptly, and many pioneering ideas remain dormant until later researchers recognize their value (Van Raan, 2004).

In the digital era, scientific results are published at an astonishing rate (Landhuis, 2016), with the number of scientific articles published annually more than tripling over the past two decades, surpassing six million papers in 2023 (www.dimensions.ai). The field of ecology is no exception to this trend (Pautasso, 2012), as researchers struggle to keep up with the ever-growing influx of new literature (Courchamp & Bradshaw, 2018). As a result, readers must be more selective in what they consume (Mabe & Amin, 2002), while writers may adapt their language to capture attention (Weinberger et al., 2015; França & Monserrat, 2019; Mammola, 2020). Further, journals may reinforce this trend by requiring authors to emphasize the novelty of their publications. As readers striving to keep up with the relentless production of ecological literature, we sensed that an increasing number of papers are filled with terms that, in various ways, highlight the novelty of the research. Here, we explore the question: Is this trend real or merely perceived?

We analyzed the relative use (i.e. frequency) of novelty and confirmatory terms in ecological publications over a twenty-year period. We developed a dual-hypothesis testing framework (Fig. 1). If ecological research is primarily confirmatory, we would expect a consistently higher relative use of confirmatory terms than novelty terms (H1; Fig. 1A,C). Conversely, if the pressure to stand out in the “research crowd” influences authors’ writing and journal publishing practices, we should observe a significant increase in the relative use of novelty terms over time (H2; Fig. 1B,C).

**Figure 1.**
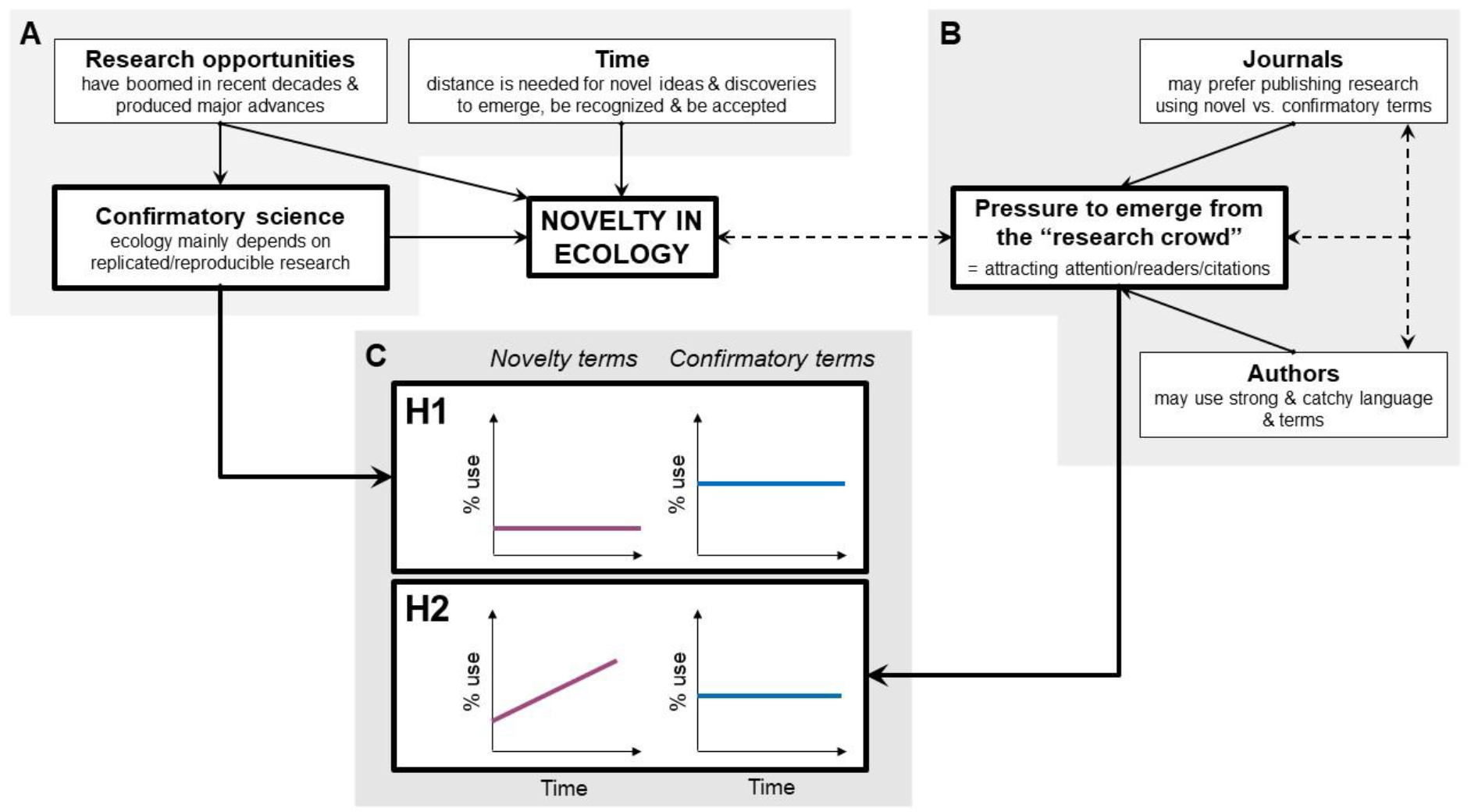
Schematic of the dual-hypothesis framework. The confirmatory nature of ecological research (A) contrasts with the pressure on authors and journals to stand out in an increasingly crowded research landscape (B), leading to two distinct scenarios (C). Solid arrows indicate putative direct relationships between components, while dashed arrows represent plausible interactions or synergies that, in turn, shape the hypothesized temporal patterns in the use of novelty and confirmatory terms.

Additionally, we conducted a scientometrics analysis to examine whether relationships exist between the use of novelty or confirmatory terms and (i) the Impact Factor (Journal Impact Factor) of the journal in which a paper was published or (ii) the number of citations a paper received. A relationship with Journal Impact Factor would suggest a journal’s tendency to either favor (positive relationship) or discourage (negative relationship) papers using these terms. In a more subtle way, this pattern may also reflect the influence of editorial and reviewer preferences, shaped by the perceived prestige of journals, rather than any intrinsic characteristic of the journals themselves. A relationship with citation count would indicate whether readers are more (positive relationship) or less (negative relationship) likely to cite papers containing either type of term.

## Dataset and statistical analyses

We used a dataset of 52,236 papers published between 1997 (year in which Journal Impact Factor was introduced) and 2017 in 17 representative ecological journals (Mammola et al., 2021) (Table S1) – these constituting ∼20% of all ecological journals listed in the Web of Science in 1997, and ∼11% of those listed in 2017, and covering most of the Journal Impact Factor range in ecology (e.g. 1.3-10.8 for the year 2023). We examined the frequency of appearance (use/non-use) of a set of selected novelty terms (“breakthrough”, “groundbreaking”, “innovated”, “innovation”, “innovative”, “new”, “newly”, “novel”, “novelty”) and confirmatory terms (“confirm”, “confirmatory”, “replicability”, “replicate”, “replicated”, “replication”, “reproducibility”) over time in paper abstracts (i.e. scoring a “use” for at least one novelty/confirmatory word). We focused on abstracts because they reflect the overall writing style of articles (Plavén-Sigray et al., 2017), while representing the lark mirror to capture the attention of readers (Martínez & Mammola, 2021).

We used regression-like analyses (Zuur & Ieno, 2016) to examine whether the use of novelty or confirmatory terms has increased over the studied period across all papers and journals (N = 52,236). Specifically, we ran two generalized linear mixed models to test the relationship between the use of confirmatory and novelty terms and publication year, with ‘journal’ included as a random-intercept factor, assuming that abstracts from the same journal share more similar writing features than those from different journals. Given the binary nature of the dependent variable (0 = non-use of the term; 1 = use of the term in each paper), we specified a Bernoulli-family data distribution and a complementary log-log link function to account for the unbalanced distribution of zeros and ones. To provide a visual summary of the temporal trend, we plotted the frequency of term usage as the percentage of papers using novelty or confirmatory terms per year—both in aggregate (Fig. 2) and for individual journals (Fig. 3).

**Figure 2.**
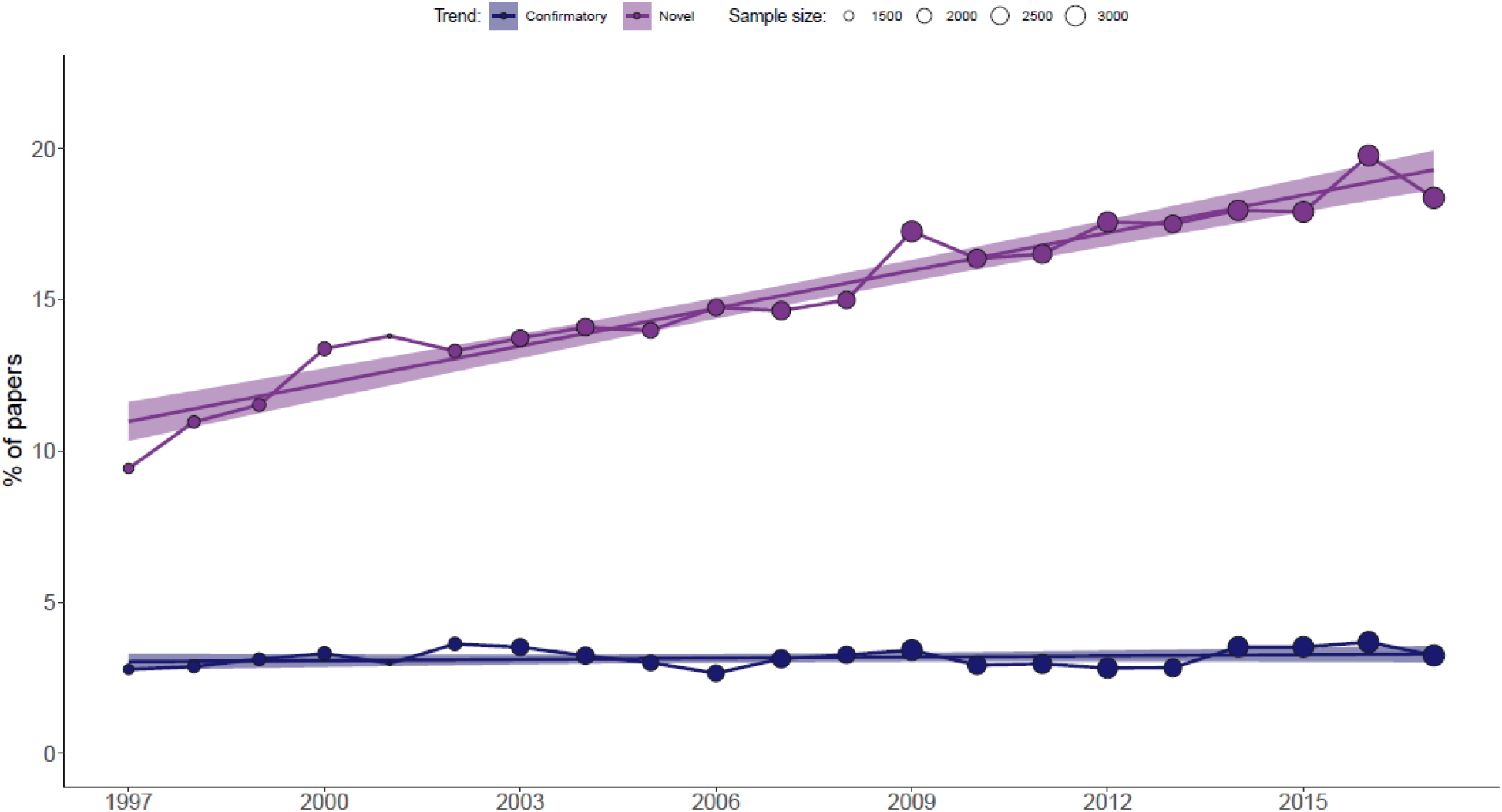
Increasing use of novelty terms in ecological abstracts. Temporal trends in the relative use (i.e. annual frequency [%]) of novelty and confirmatory terms across 17 selected ecological journals (Table S1). Dot size represents the number of articles published each year. Regression lines and confidence intervals are included for visual clarity, based on a linear model fitted through the data.

**Figure 3.**
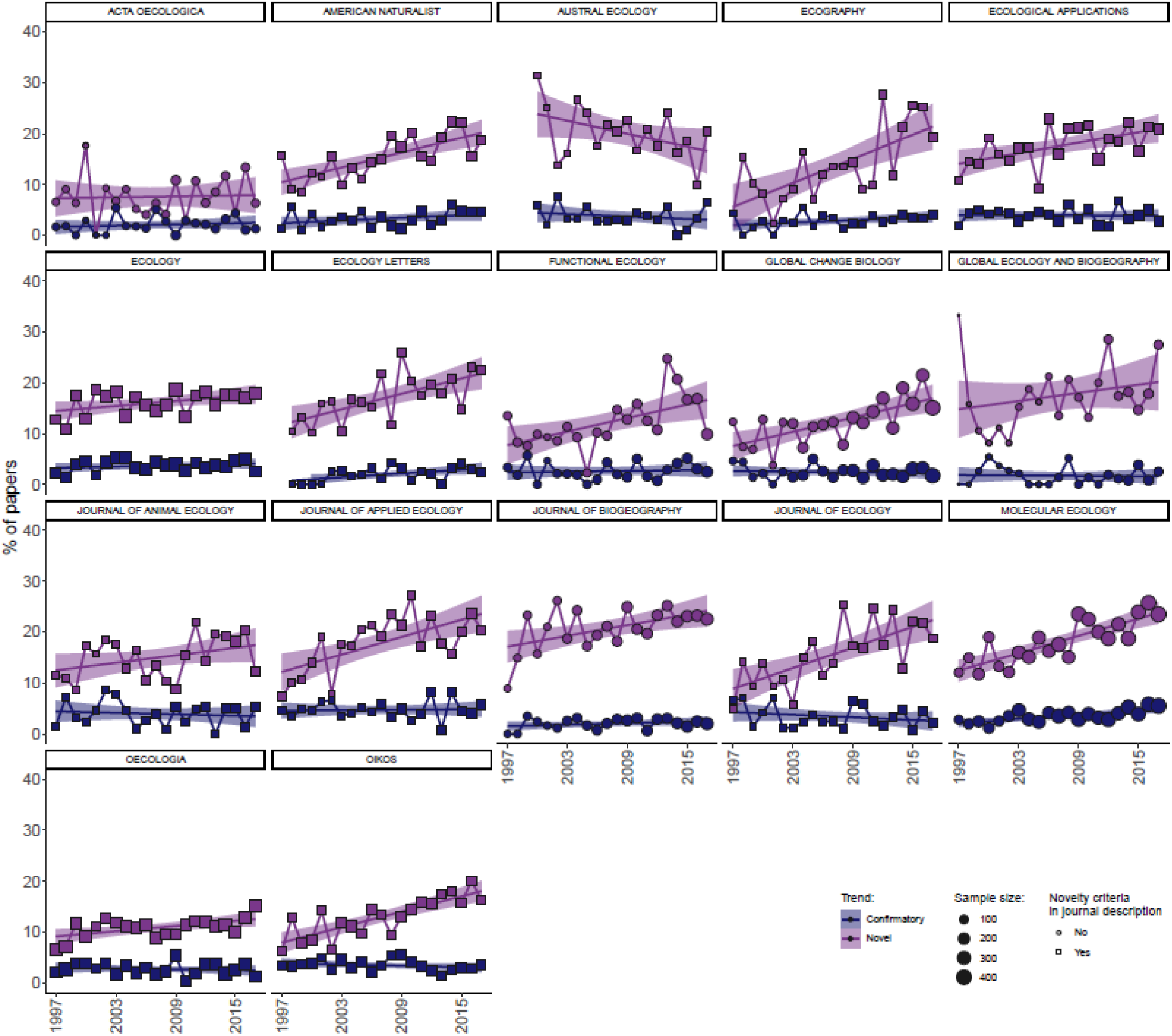
The trend of increasing use of novelty terms in ecological abstracts is consistent across all but one journal. Temporal trends in the relative use (i.e. annual frequency [%]) of novelty and confirmatory terms for each of the 17 selected ecological journals. Symbols indicate whether novelty is a criterion mentioned in the journal description (Table S1), and their size corresponds to the number of articles published each year. Regression lines and confidence intervals are included for visual clarity.

Next, we used a generalized linear mixed model to test whether the number of citations (response variable) is related to the relative use of novelty and confirmatory terms (fixed effects). We also included abstract length (word count) and publication year as covariates to control their potential influence on citations, and we treated ‘journal’ as a random-intercept factor. Since citations are count data, we initially specified a Poisson-family distribution. However, the Poisson model was highly over-dispersed (dispersion ratio = 96.5, Pearson’s χ^2^ = 5040868.5, p < 0.001), so we switched to a negative binomial distribution. To examine whether the use of novelty and confirmatory terms is related to Journal Impact Factor, we ran a linear model with the same fixed effects as in the citation model. Each paper was assigned the Journal Impact Factor corresponding to its year of publication. Here, we did not include ‘journal’ as a random effect, as it is inherently tied to Journal Impact Factor. It must be pointed out that, technically, a Gaussian distribution may not be the most appropriate choice in this instance (as Journal Impact Factor values cannot assume negative values). However, given that the linear model validation yielded satisfactory results, we opted to retain the simpler approach rather than adopting a more complex distribution (e.g. Gamma).

We ran all the analyses in R version 4.3.0 (R Core Team, 2023), using the package glmmTMB version1.1.7 for regression analyses (Brooks et al., 2017), performance version 0.9–7 for model validation (Lüdecke et al., 2021), and ggplot2 version 3.5.1 for plotting (Wickham et al., 2016).

## The growing use of novelty terms in ecology

Across all journals, the relative use of novelty terms in paper abstracts doubled over the study period, increasing from ∼10% in 1997 to ∼20% in 2017 (Fig. 2). Logistic regression analyses confirmed that the likelihood of an article using novelty terms was higher in recent years (*Log-Risk* ± SE: 0.16 ± 0.01, *z* = 14.03, *p* < 0.001; Conditional *R*^*2*^ = 0.05, Marginal *R*^*2*^ = 0.02). In contrast, we found no clear trend for confirmatory terms, whose relative use remained steady at around 3% (Fig. 2). The probability of an article using confirmatory terms also remained stable over the study period (*Log-Risk* ± SE: 0.04 ± 0.02, *z* = 1.54, *p* = 0.125; Conditional *R*^*2*^ = 0.03, Marginal *R*^*2*^ = 0.01). This overall pattern for novelty and confirmatory terms was similar across all journals, except for *Austral Ecology*, which—*nomen omen*—showed the opposite trend, with the use of novelty terms declining over time (Fig. 3).

The use of novelty terms was positively associated with both the number of citations and Journal Impact Factor, whereas no such relationships were found for confirmatory terms (Fig. 4). Abstract length (number of words) was positively associated with the number of citations and negatively with Journal Impact Factor, while publication year was negatively related to the number of citations (i.e. more recent papers receive fewer citations than older ones) and positively with Journal Impact Factor. The unexplained variance suggests that several other factors, not accounted for in this analysis, are likely influencing article impact—something that is well-documented in the “science of science” literature (e.g. Tahamtan et al., 2016, 2019; Mammola et al., 2022).

**Figure 4.**
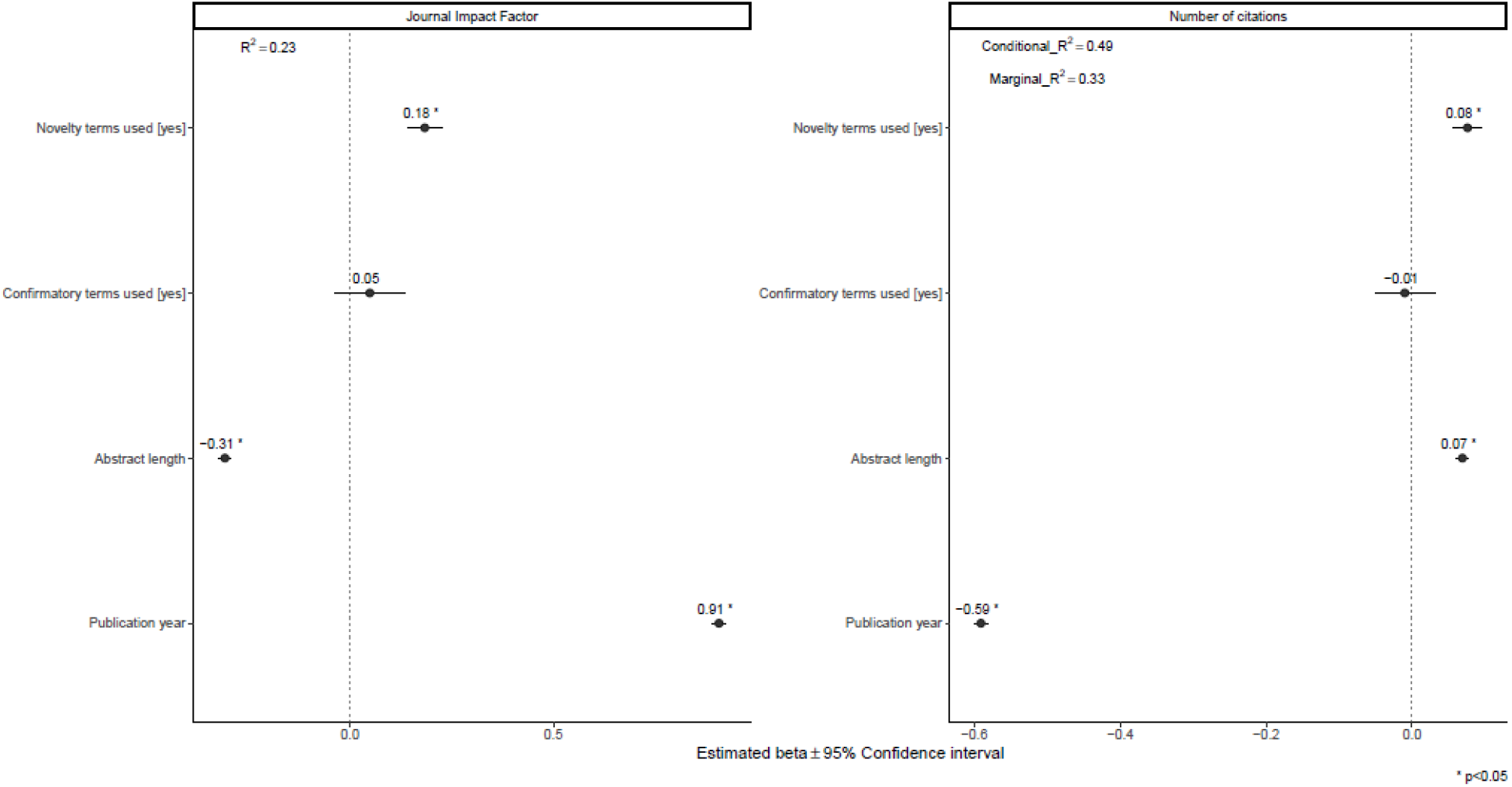
Publication impact is tightly associated with the use of novelty terms. Forest plots summarize the estimated parameters of regression models testing the relationship between novelty and confirmatory terms, abstract length (number of words), and publication year on the Journal Impact Factor (left panel; based on a linear model) and the number of citations (right panel; based on a generalized linear mixed model). Bars represent 95% confidence intervals. Variance explained is reported as both conditional *R*^*2*^ (random + fixed effects) and marginal *R*^*2*^ (explained by fixed factors alone). Asterisks (*) indicate significant effects (*α* = 0.05).

## What could be behind the rising trend of novelty terms?

We found strong evidence supporting our perception that more and more papers are using novelty terms, while confirmatory terms showed no obvious temporal patterns and were generally much less used by researchers over the studied 20-year timespan (Fig. 2, Fig. 3). Concurrently, the use of novelty terms tended to attract more citations and was associated with journals having higher Journal Impact Factors compared to the use of confirmatory terms (Fig. 4). As a result, we rejected H1 of our dual-hypothesis framework, while H2 received striking support (Fig. 1). The increasing use of novelty terms was confirmed across all our analyses, emerging across all journals (Fig. 2), as well as within individual journals (Fig. 3). The only exception was the Australian journal *Austral Ecology*, which exhibited a temporal decline in the relative use of novelty terms, for which we do not have a plausible explanation for this anomalous “down-under” pattern. Taken together, these findings support the idea that the pressure to stand out from the “research crowd” felt by both researchers and journals plays a key role in the current ecological writing and publishing landscape (Fig. 1).

Still, we can only speculate about the possible causes driving the upward trend in the use of novelty terms in the last two decades, as correlation does not necessarily imply causation. Perhaps, thanks to recent conceptual developments (Dubois & Peres-Neto, 2022) and the increasing availability of data and analytical tools (e.g. Besson et al., 2022; Cardoso et al., 2020; McCallen et al., 2019; Tosa et al., 2021; Mammides & Papadopoulos, 2024), ecologists are now truly able to make groundbreaking discoveries and write novel stories at an accelerating pace. However, the history of science suggests that game-changing findings are rare and take time to be recognized (Morris, 2009; Van Raan, 2004). This view is further supported by a recent overview illustrating how papers are increasingly less likely to make scientific breakthroughs (Park et al., 2023).

We must then face an uncomfortable alternative possibility: are we, as ecologists, using a more sensationalized and novelty-driven language (either consciously or unconsciously) to increase our chances of catching readers’ attention amidst the incessant production of scientific literature (scenario depicted in Fig. 1B, C) (Weinberger et al., 2015; Doubleday & Connell, 2017; Mammola, 2020)? This speculation is supported by the positive significant relationship between the use of novelty terms, but not the use of confirmatory terms, and both number of citations and Journal Impact Factor (Fig. 4). These relationships also suggest that Journal Impact Factor could benefit from publishing papers that use novelty terms, as they are more likely to attract citations. Indeed, journals may be contributing to this trend. Among the 17 ecological journals included in our analysis, about 65% explicitly mention novelty as a criterion in their current author guidelines (Table S1). Similarly, novelty is a core requirement in pre-peer review editorial decisions for some journals (Arnqvist, 2013).

Thus, this “quest for novelty” may partly stem from the challenges faced by journals in attracting readers and citations. At the same time, more “novel” papers may tend to be published in journals with higher Journal Impact Factor, further shaping the observed patterns. In other words, such complex feedback loops between researchers and journals may therefore largely contribute to generating the spike in the use of novelty terms in ecological literature.

## Limitations of the study

A deeper mechanistic understanding of what drives these scientometrics patterns related to writing and publishing behaviors would require a closer examination of each manuscript included in this study. This step would imply reading each of the >50k papers, and perhaps even contacting corresponding authors asking for their feedback and reasons behind the choice of using or not novelty terms. We are also aware that the selection of terms and searched journals can affect the revealed patterns. However, thanks to the representativeness of the chosen ecological journals, Journal Impact Factor range, and set of selected terms, we are confident that what we have found offers a reliable picture of what has happened in the studied 20-year timespan.

## On the importance and impacts of confirmatory science and of language use in ecology

Ecology is experiencing unprecedented research opportunities worldwide. However, like any other scientific discipline, knowledge-building progresses through a lengthy and steady cumulative process, with most basic and applied research being inherently confirmatory in nature (Hoyningen-Huene, 2013). Novel ideas and discoveries may emerge in response to idiosyncrasies arising from observational or experimental studies, which also form the theoretical foundations upon which we built—and ultimately rejected—our H1. Nevertheless, the frequency of new discoveries in ecology typically occurs at a rate of only a few per year or decade (Morris, 2009), which contrasts with the trends we observed in our study.

From a semantic and cognitive standpoint, words are not just tools for communicating our key findings to other scientists or the broader public (Feynman, 1969), but also serve as the building blocks of knowledge construction (Martínez & Mammola, 2021). We wonder whether the increasing use of sensationalized language (Mammola, 2020), where novelty may be exaggerated, could influence our thinking process at various levels. After all, understanding what is truly new is crucial—not only when writing and disseminating results but also when designing future projects and experiments. Without this clarity, we risk reinventing the wheel. We join the call to evaluate publications based on their quality, soundness, clarity, and replicability, giving less emphasis to their confirmatory or novelty (true or claimed) nature (Pautasso, 2013; Romero, 2017). Encouragingly, this approach seems to be increasingly adopted by ecological journals, especially (but not exclusively) open-access ones. Therefore, we emphasize the importance of starting a conversation about the potential root-causes and implications of this linguistic and scientometrics trend for the scientific community and science communication at large.

## Author contributions

GO conceived the research idea, with significant inputs to further develop it provided by SM, AM, MPB. SM gathered the data and conducted the statistical analysis. GO and SM led the writing, and all coauthors contributed to revisions.

## Acknowledgements

We thank François Munoz (recommender at PCI Ecology), François Massol and Matthias Grenié for providing very constructive feedback during the revision process.

## Data accessibility statement

Data supporting this study is available in Figshare: https://doi.org/10.6084/m9.figshare.12941639.v1.

The analytical pipeline to reproduce the analyses is also available in GitHub: https://github.com/StefanoMammola/Ottaviani_et_al.

## Artificial Intelligence (AI) declaration

No AI technologies have been used.

---

**Table S1.** The 17 journals selected for the analysis and sample size (readapted from Mammola et al., 2021).

| Journal name | Initial year | Timespan selected | N of primary research articles used | Use and requirement of novelty terms in journal description* |
| --- | --- | --- | --- | --- |
| Acta Oecologica | 1983 | 1997–2017 | 1,408 | No |
| American Naturalist | 1867 | 1997–2017 | 2,852 | Yes |
| Austral Ecology | 2000 | 2000–2017 | 1,434 | Yes |
| Ecography | 1978 | 1997–2017 | 1,743 | Yes |
| Ecological Applications | 1991 | 1997–2017 | 3,051 | Yes |
| Ecology | 1920 | 1997–2017 | 5,505 | Yes |
| Ecology Letters | 1998 | 1998–2017 | 2,098 | Yes |
| Functional Ecology | 1987 | 1997–2017 | 2,326 | No |
| Global Change Biology | 1995 | 1997–2017 | 3,937 | No |
| Global Ecology and Biogeography | 1993 | 1997–2017 | 1,377 | No |
| Journal of Animal Ecology | 1932 | 1997–2017 | 2,250 | Yes |
| Journal of Applied Ecology | 1964 | 1997–2017 | 2,407 | Yes |
| Journal of Biogeography | 1974 | 1997–2017 | 2,852 | No |
| Journal of Ecology | 1913 | 1997–2017 | 2,170 | Yes |
| Molecular Ecology | 1992 | 1997–2017 | 6,209 | No |
| Oecologia | 1968 | 1997–2017 | 5,446 | Yes |
| Oikos | 1949 | 1997–2017 | 3,812 | Yes |

\* Novelty terms considered in the journal description (i.e. scope and authors' guidelines; search conducted in 2021) are the same as of the paper abstract search.

